# Plug-and-play assembly of biodegradable ionizable lipids for potent mRNA delivery and gene editing *in vivo*

**DOI:** 10.1101/2025.02.25.640222

**Authors:** Xuexiang Han, Ying Xu, Adele Ricciardi, Junchao Xu, Rohan Palanki, Vivek Chowdhary, Lulu Xue, Ningqiang Gong, Mohamad-Gabriel Alameh, William H. Peranteau, James M. Wilson, Drew Weissman, Michael J. Mitchell

**Affiliations:** Department of Bioengineering, University of Pennsylvania, Philadelphia, PA 19104, USA; Key Laboratory of RNA Innovation, Science and Engineering, CAS Center for Excellence in Molecular Cell Science, Shanghai Institute of Biochemistry and Cell Biology, Chinese Academy of Sciences, University of Chinese Academy of Sciences, Shanghai 200031, China; Department of Medicine, University of Pennsylvania, Philadelphia, PA 19104, USA; Penn Institute for RNA Innovation, Perelman School of Medicine, University of Pennsylvania, Philadelphia, PA 19104, USA; School of Pharmacy, East China Normal University, Shanghai 200062, China; Division of Pediatric General, Thoracic, and Fetal Surgery, The Center for Fetal Research, Children’s Hospital of Philadelphia, Philadelphia, PA 19104, USA; Gene Therapy Program, Perelman School of Medicine, University of Pennsylvania, Philadelphia, PA 19104, USA; Abramson Cancer Center, Perelman School of Medicine, University of Pennsylvania, Philadelphia, PA 19104, USA; Institute for Immunology, Perelman School of Medicine, University of Pennsylvania, Philadelphia, PA 19104, USA; Cardiovascular Institute, Perelman School of Medicine, University of Pennsylvania, Philadelphia, PA 19104, USA; Institute for Regenerative Medicine, Perelman School of Medicine, University of Pennsylvania, Philadelphia, PA 19104, USA

## Abstract

mRNA-based gene editing therapeutics offer the potential to permanently cure diseases but are hindered by suboptimal delivery platforms. Here, we devise a robust combinatorial chemistry for plug-and-play assembly of diverse biodegradable ionizable lipids and identify a lead candidate that produces superior lipid nanoparticles for various gene editing tools delivery *in vivo*. Our study highlights the utility of this synthetic approach and the generality of this platform for potent *in vivo* gene editing.

## Main

Clustered regularly interspaced short palindromic repeat (CRISPR)-based gene editing technology provides a precise molecular mechanism for genome manipulation^1^, which has been exploited to develop various therapeutic tools to treat diseases^2^, especially genetic disorders. Compared to other therapeutic modalities, gene editing has the potential to cure diseases permanently with only a single treatment. However, a key challenge for gene editing therapeutics is their safe and efficient delivery *in vivo*. While viral vectors, such as adeno-associated virus, show high transfection activity, their *in vivo* gene editing application is restricted by strong immunogenicity, limited packaging capacity, persistent nuclease expression, potential insertional mutagenesis, and high production cost^3^.

These limitations can be overcome by lipid nanoparticles (LNPs), the most clinically advanced nonviral vector^4^. Through co-delivery of mRNA-encoded CRISPR-associated nuclease and single guide RNA (sgRNA) specific for a target locus in the genome, LNPs have successfully achieved *in vivo* gene editing in both preclinical and clinical studies^5, 6^. However, compared to mRNA vaccines, mRNA-encoded CRISPR therapeutics require up to 1,000-fold-higher mRNA dosage (or protein expression) to reach a therapeutic threshold^7^, which raises higher requirements for the potency and safety of LNPs. Therefore, there is immense interest in developing next-generation ionizable lipids (ILs) – a critical component of LNPs that governs the efficacy and biocompatibility. Moreover, new methods for simple, robust and scalable synthesis of these ILs are valuable for the practical application.

Herein, we developed a plug-and-play assembly strategy for the combinatorial synthesis of biodegradable ILs. Compared to benchmark ILs, our lead IL – synthesized in one step from an amine and a dialkyl maleate – demonstrates superior performance in delivering various CRISPR mRNA therapeutics in wild-type and transgenic mice. Our study highlights the utility of this combinatorial chemistry as well as the broad capability of the lead IL for *in vivo* gene editing.

In this study, we devised a two-component combinatorial chemistry for “plug-and-play” assembly of biodegradable ILs between amines/thiols and dialkyl maleates (**Fig. 1a**). This “click-like” Michael addition reaction has many advantages, such as easily accessible building blocks, a simple experimental procedure, avoidance of solvent and catalyst, 100% atom economy, and a high product yield^8^. Notably, compared with previously reported Michael addition reaction for IL synthesis that uses alkyl acrylate^4, 9, 10^, the use of dialkyl maleate in our scheme has several benefits: first, dialkyl maleate is more reactive than alkyl acrylate, due to the two electron-withdrawing groups to the double bond^8^; second, unlike alkyl acrylate, that uncontrollably produces a mixture of both mono- and bis-adduct, dialkyl maleate is highly selective towards the mono-adduct for primary amines due to the high steric hindrance^8^, leading to less by-products; third, ILs with asymmetric tails can be easily prepared from asymmetric dialkyl maleate (**Fig. 1a** and **Supplementary Fig. 1**), which expands the structural diversity of ILs^11^.

**Fig. 1.**
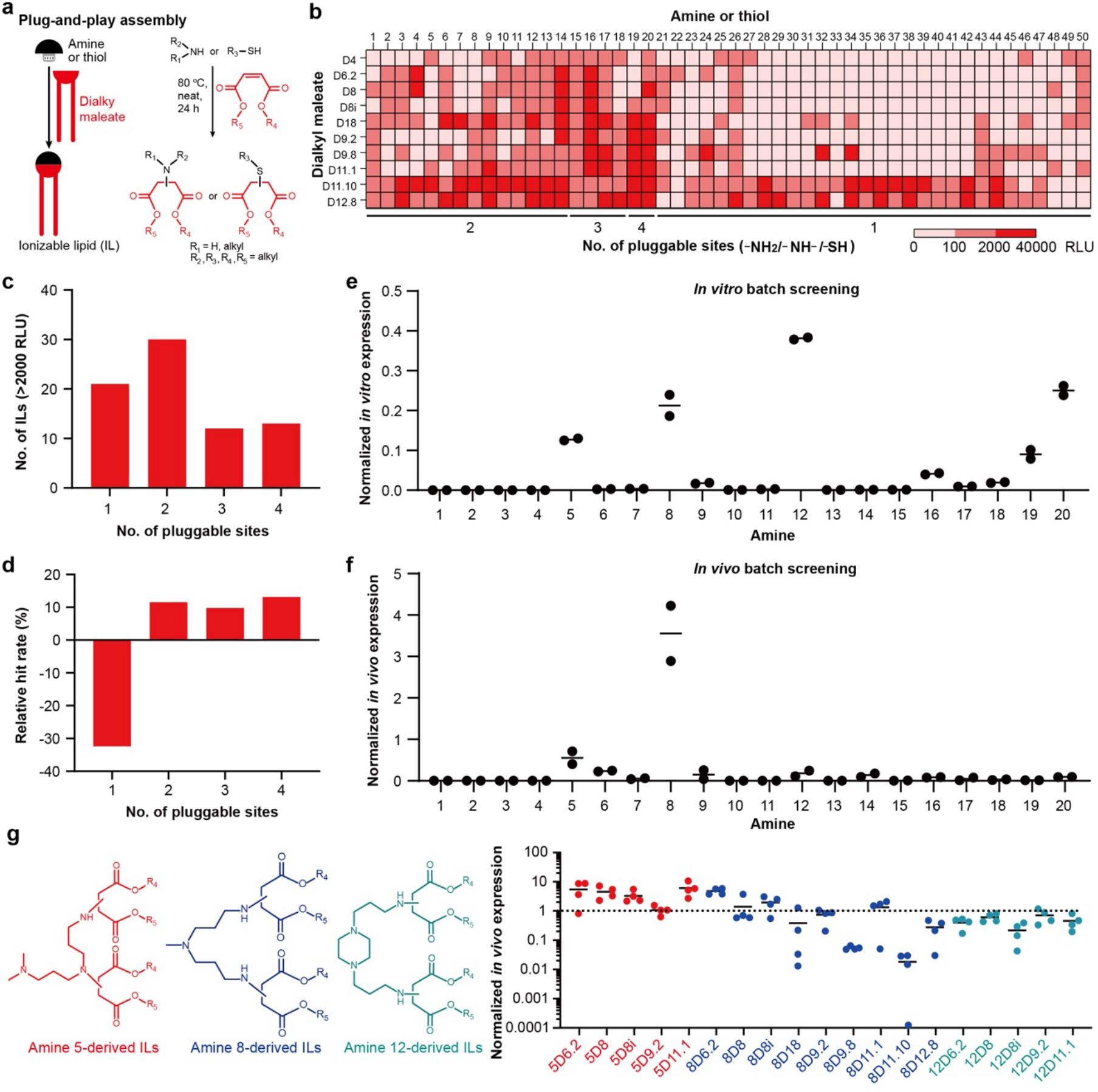
Plug-and-play assembly of ILs and *in vitro* and *in vivo* screening. **a** Plug-and-play assembly of biodegradable ILs through Michael addition between amines/thiols and dialkyl maleates. **b** Heat map of *in vitro* luciferase expression. Data are presented as mean (*n* = 2 independent biological replicates, initial screening). HepG2 cells were treated with mLuc-loaded LNPs at an mRNA dose of 3 ng/well for 24 h. RLU, relative light unit. Untreated cells typically exhibited <100 RLU. **c** Distribution of ILs with *in vitro* luciferase expression >2000 RLU based on the number of pluggable sites. **d** Analysis of relative hit rate (percentage in the group with luciferase expression >2000 RLU – percentage in library) based on the number of pluggable sites. **e** *In vitro* luciferase expression for batch LNP screening. mLuc-LNPs with the same amine structure were pooled and used to treat HepG2 cells at an mRNA dose of 15 ng/well for 24 h. Data are normalized to C12-200 LNP and presented as mean (*n* = 2 independent biological replicates, initial screening). **f** *In vivo* luciferase expression for batch LNP screening. mLuc-LNPs with the same amine structure were pooled and injected into mice at an mRNA dose of 0.1 mg/kg. Data are normalized to C12-200 LNP and presented as mean (*n* = 2 independent biological replicates, initial screening). **g** *In vivo* luciferase expression for individual LNP screening. Selected ILs derived from amine 5, 8 and 12 were formulated into mLuc-LNPs individually, which were injected into mice at an mRNA dose of 0.1 mg/kg. Data are normalized to C12-200 LNP and presented as mean (*n* = 4 independent biological replicates).

We combinatorially synthesized 500 ILs using 50 amines/thiols and 10 dialkyl maleates (**Fig. 1b** and **Supplementary Fig. 1**). Notably, amines/thiols 1-50 are classified based on the number of reaction sites (i.e. –NH_2_/–NH–/–SH) with dialkyl maleate. Symmetric dialkyl maleates (i.e. D4, D6.2, D8 and D8i) are commercially available, while asymmetric ones (i.e. D18, D9.2, D9.8, D11.1 D11.10 and D12.8), that are variable in length, branching, and saturation, were synthesized through the one-step esterification reaction (**Supplementary Fig. 2**). Each IL was named according to the specific combination of the amine/thiol and the dialkyl maleate used in its synthesis. These ILs were individually formulated into mLuc-LNPs along with other lipid excipients and firefly luciferase mRNA (mLuc), which were then subjected to high-throughput *in vitro* screening. HepG2 cells (a human hepatocellular carcinoma cell line) were treated with mLuc-LNPs at a low mRNA dose (3 ng/well) to avoid cytotoxicity, and luciferase expression was measured after 24 hours (**Fig. 1b**). The *in vitro* transfection efficiency of ILs assembled from amine 1-20 with two to four pluggable sites was greatly affected by amine structure. Overall, amines bearing multiple pluggable sites (e.g. amine 20) afforded efficacious ILs. In contrast, tail structure profoundly impacted the transfection efficiency of ILs assembled from amine 21-47 with only one pluggable site. Generally, only dialkyl maleate with a long tail and a branch (e.g. D12.8) generated efficacious ILs. In contrast, thiols (e.g., thiolated amine) 48-50 with one pluggable site resulted in suboptimal ILs irrelevant of tail structure.

**Fig. 2.**
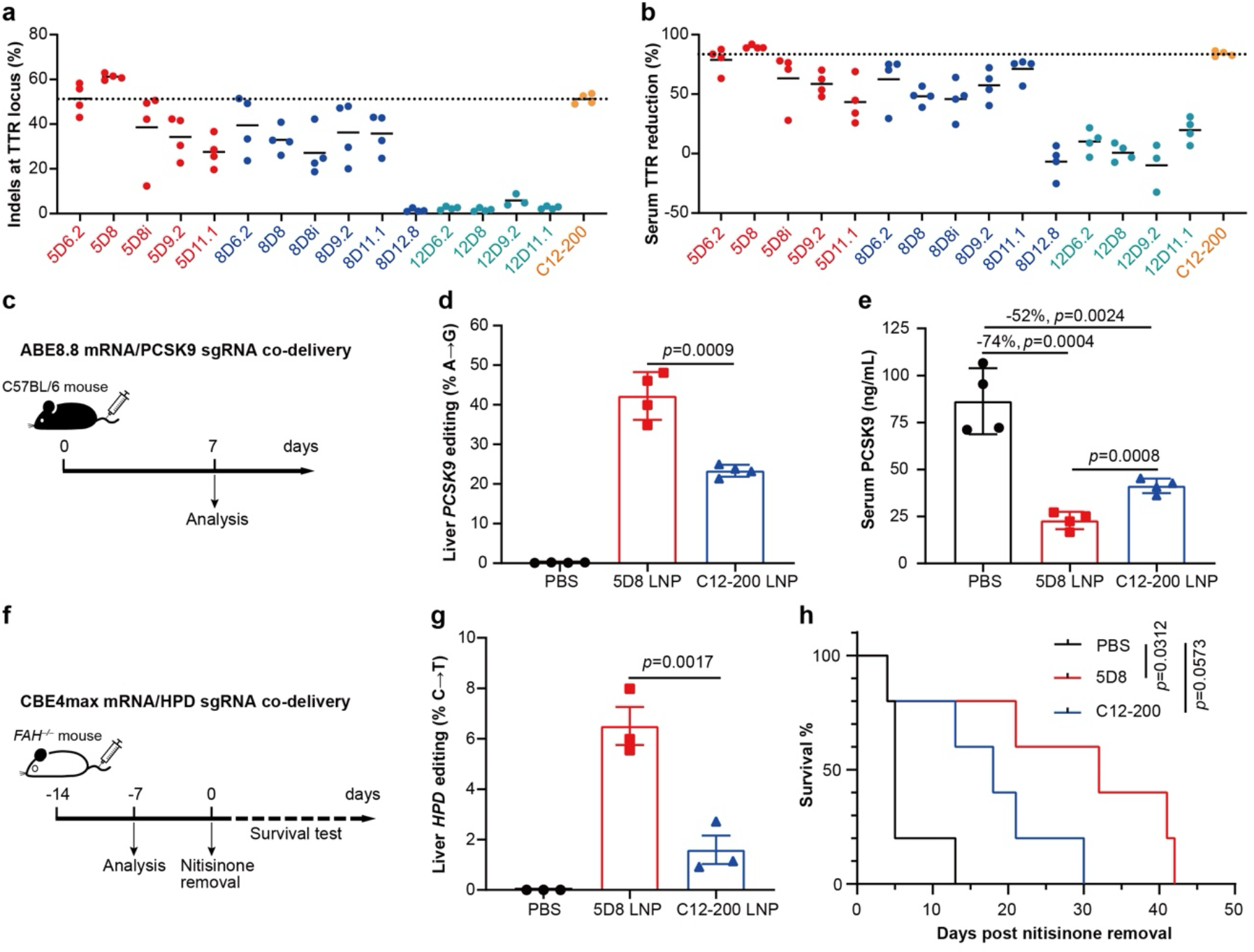
LNP-mediated *in vivo* gene editing. **a, b** TTR gene editing and serum TTR reduction. LNPs encapsulating Cas9 mRNA/TTR sgRNA (4:1, wt:wt) were i.v. injected into C57BL/6 mice at a total RNA dose of 1 mg/kg. On day 7, DNA was extracted from the liver to determine on-target indel frequency by next-generation sequencing (**a**, *n* = 3-4 independent biological replicates), and serum was collected for ELISA analysis of TTR (**b**, *n* = 3-4 independent biological replicates). Data are presented as mean. **c**–**e** PCSK9 base editing. LNPs encapsulating ABE8.8 mRNA/PCSK9 sgRNA (4:1, wt:wt) were i.v. injected into C57BL/6 mice at a total RNA dose of 0.75 mg/kg (**c**). On day 7, DNA was extracted from the liver to determine on-target editing frequency by next-generation sequencing (**d**, *n* = 4 independent biological replicates), and serum was collected for ELISA analysis of PCSK9 (**e**, *n* = 4 independent biological replicates). Data are presented as mean ± SD. Statistical significance was evaluated by a one-way ANOVA with Tukey’s correction. **f**–**h** HPD base editing. LNPs encapsulating CBE4max mRNA/HPD sgRNA (4:1, wt:wt) were i.v. injected into *FAH*^−/−^ mice at a total RNA dose of 0.6 mg/kg (**f**). Seven days later, DNA was extracted from the liver to determine on-target editing frequency by next-generation sequencing (**g**, *n* = 3 independent biological replicates). Data are presented as mean ± SD. Statistical significance was evaluated by a one-way ANOVA with Tukey’s correction. Two weeks later, nitisinone was removed and the survival of *FAH*^−/−^ mice was monitored (**h**, *n* = 5 independent biological replicates). Statistical significance was evaluated using Survival Curve with Log-rank (Mantel-Cox) test.

Through this screen, we identified 76 ILs that mediated *in vitro* mRNA delivery with luciferase expression >2000 relative light unit (RLU, **Fig. 1b, c**). Notably, none of these ILs were assembled from dialkyl maleate D4, presumably due to its short hydrocarbon chains and low lipophilicity (**Supplementary Fig. 1**). For amine/thiol with one to four pluggable sites, relative hit rates (percentage in the group with luciferase expression >2000 RLU – percentage in library) were calculated to be −32.4, 11.5, 9.8 and 13.1, respectively (**Fig. 1d**). These results suggest that amine/thiol 21-50 with one pluggable site is inferior in affording efficacious ILs compared to other counterparts. Thus, we focused on the 200 ILs assembled from amine 1-20 in the following studies.

To accelerate the identification of optimal amines and reduce the usage of animals, we utilized a batch screening method^12, 13^. mLuc-LNPs with the same amine structure were pooled into 20 batches (10 LNPs per batch), which were then used to treat HepG2 cells or intravenously (i.v.) injected into mice. Amine 12 was identified as the most effective based on our *in vitro* results, followed by amine 20, 8 and 5 (**Fig. 1e**). However, none of these pooled LNPs reached the transfection efficiency of C12-200 LNP, a potent benchmark for the delivery of mRNA therapeutics and gene editors^14^. Interestingly, *in vivo* batch screening results showed that amine 8 and 5 were the best performers and achieved higher (3.6-fold) and slightly lower (0.6-fold) transfection efficiency relative to C12-200 LNP (**Fig. 1f** and **Supplementary Fig. 3**), respectively.

Taking the above results into consideration, we next chose all amine 8-derived ILs, except 8D4, for individual screening in mice. For rigid comparison, these ILs were purified (**Supplementary Fig. 4**), and formulated into mLuc-loaded LNPs using a microfluidic device. *In vivo* results demonstrated that LNPs formulated with 8D6.2, 8D8, 8D8i, 8D9.2 or 8D11.1 achieved comparable or higher transfection efficiency than C12-200 LNP (**Fig. 1g**). Notably, ILs with increased tail and branch length tended to show decreased transfection activity (e.g. 8D8 *vs* 8D18; 8D9.2 *vs* 8D9.8). These results are in line with previous studies, where potent ILs often contain multiple short tails^9, 10, 15^. Following this discovery, we further tested amine 5-derived five ILs with multiple short tails (**Fig. 1g**), all of which were comparable or superior to C12-200. Due to the strong *in vitro* transfection activity, amine 12-drived five ILs with multiple short tails were also tested *in vivo*, but all of them were slightly inferior to C12-200.

Inspired by the ease of synthesis and highly efficient mRNA delivery *in vivo*, we tested the utility of these ILs to deliver CRISPR mRNA therapeutics for *in vivo* gene editing. Ten superior ILs from amine 5 and 8 were screened for liver genome editing by co-delivering Cas9 mRNA and sgRNA targeting the transthyretin (*TTR*) gene in mice (**Fig. 2a, b**). Five inferior ILs in our initial screen were also tested for comparison. Encouragingly, all ten superior ILs achieved more than 27% on-target editing efficiency. The lead 5D8-formulated LNP achieved the highest on-target editing efficiency (~61%), which was higher than C12-200 LNP (~51%, **Fig. 2a**) and other industry benchmark LNPs, including SM-102 LNP (~20%), MC3 LNP (~5%), and LP-01 LNP (~3%) (**Supplementary Fig. 5**). Correspondingly, serum TTR protein was reduced by ~90% in mice treated with 5D8 LNP (**Fig. 2b**). In contrast, all inferior ILs barely facilitated liver gene editing and serum TTR reduction. Therefore, 5D8 (characterization data in **Supplementary Figs. 6**-**9** and **Supplementary Table 1**) was identified as the top-performing ionizable lipid for CRISPR gene editing. Of note, 5D8 LNP is also highly potent for siRNA delivery, achieving ~100% reduction of serum TTR after delivery of TTR siRNA at a very low dose (0.05 mg/kg, **Supplementary Figs. 10**), which is superior to industry benchmark MC3 LNP at the same experimental conditions^16^. One possible reason for the high potency of 5D8 LNP could be its enrichment of apolipoprotein E on the surface (**Supplementary Figs. 11**), which is implicated in the active targeting of hepatocytes^17^.

We then analyzed the relationship of mLuc delivery efficiency, gene editing efficiency and serum TTR reduction (**Supplementary Fig. 12**). There was a moderate positive correlation between mLuc delivery efficiency and gene editing efficiency or serum TTR reduction, suggesting that the potency of LNPs for mLuc delivery generally predict gene editing outcome. Unsurprisingly, gene editing efficiency correlated well with serum TTR reduction^5^. We further examined the hepatotoxicity of 5D8 LNP following *in vivo* gene editing (**Supplementary Fig. 13**). There was no observable increase of alanine transaminase (ALT) or aspartate aminotransferase (AST) level following 5D8 LNP treatment. In contrast, C12-200 LNP treatment resulted in a slight elevation of both ALT and AST levels. Taken together, 5D8 is a superior ionizable lipid that enables potent mRNA delivery and gene editing with a favorable safety profile.

We next investigated whether 5D8 LNP could serve as a universal delivery platform for other mRNA-based gene editing tools, such as CRISPR base editors. Unlike CRISPR/Cas9-based gene editors, base editors enable precise and efficient base changes without the introduction of double stranded DNA breaks. Base editing can be used to permanently turn off disease-associated genes or correct pathogenic point mutations to treat single-nucleotide genetic diseases^18^. We tested the ability of 5D8 LNP to deliver adenine base editors (ABEs), which convert A•T base pairs to G•C base pairs. Mice were i.v. injected with LNPs encapsulating mRNA encoding an ABE version 8.8m (ABE8.8) and a sgRNA targeting the *PCSK9* (proprotein convertase subtilisin/kexin type 9) gene, a well-validated therapeutic target for the treatment of atherosclerotic cardiovascular disease (**Fig. 2c**)^19, 20^. 5D8 LNP induced ~42% liver *PCSK9* base-editing efficiency with a concomitant ~74% reduction of PCSK9 serum protein (**Fig. 2d, e**). In contrast, C12-200 LNP resulted in significantly lower *PCSK9* editing efficiency (~23%) and less serum PCSK9 reduction (~52%) at the same dose. These results demonstrate the promise of 5D8 LNP for *in vivo* base editing.

Finally, we explored the potential of 5D8 LNP for gene editing therapy in mice with a genetic disease. Hereditary tyrosinaemia type 1 (HT1) results from a loss of function mutation in *FAH* (fumarylacetoacetate hydrolase) gene, blocking the tyrosine catabolic pathway. Pharmacological inhibition of the upstream HPD (4-hydroxyphenylpyruvic acid dioxygenase) enzyme with nitisinone or knockout of *HPD* gene by base editing prevents the build-up of toxic metabolites and lethal liver failure^21^. LNPs comprising mRNA encoding a cytosine base editor version 4max (CBE4max) and a sgRNA targeting the *HPD* gene to introduce a C → T nonsense mutation were formulated and i.v. administered into adult *FAH*^−/−^ mice. 5D8 LNP resulted in ~6.5% on-target editing, which was significantly higher than that (~1.6%) achieved by C12-200 LNP. Consequently, mice receiving 5D8 LNP had a greater median survival time after nitisinone withdrawal (32 days), in comparison with PBS-treated group (5 days) and C12-200 LNP-treated group (18 days). Taken together, these results support that 5D8 LNP is a universal platform for efficient gene editing applications.

In summary, we developed a simple and robust combinatorial chemistry for the plug-and-play assembly of biodegradable ILs. Amine 5- and 8-derived ILs with multiple short tails showed superior *in vivo* mRNA delivery capability, among which 5D8 was identified as the lead candidate for liver gene editing with good tolerability. At clinically relevant doses, 5D8 LNP achieved higher gene editing efficiency or base editing efficiency than benchmark LNPs in wild-type and transgenic mice. Future efforts to improve the editing efficiency by optimizing cargos (e.g. sequences, and chemical modifications) are on-going. Nevertheless, this study demonstrates the great value of this plug-and-play assembly strategy and 5D8 LNP as a general platform for potent *in vivo* gene editing.

## Materials and Methods Materials

Amines, aliphatic alcohols, N,N’-dicyclohexylcarbodiimide (DCC), 4-dimethylaminopyridine (DMAP) and were purchased from Sigma Aldrich, Tokyo Chemical Industry, Ambeed and AstaTech. Dibutyl maleate (D4), Bis(2-ethylhexyl) maleate (D6.2), dioctyl maleate (D8), Bis(6-methylheptyl) maleate (D8i) and (Z)-4-((2-ethylhexyl)oxy)-4-oxobut-2-enoic acid were obtained from Ambeed and AstaTech. 1,2-dioleoyl-sn-glycero-3-phosphoethanolamine (DOPE), 1,2-distearoyl-sn-glycero-3-phosphocholine (DSPC), 1,2-dimyristoyl-rac-glycero-3-methoxypolyethylene glycol-2000 (DMG-PEG 2000) and cholesterol were obtained from Avanti Polar Lipids. Ionizable lipids C12-200, MC3 and SM-102 were purchased from MedChem Express. LP-01 was purchased from Cayman Chemical. Nucleoside-modified luciferase mRNA (5moU) and Cas9 mRNA (5moU) were bought from TriLink. TTR siRNAs (#NM_013697, siRNA IDs: SASI_Mm01_00076059, SASI_Mm01_00076060 and SASI_Mm01_00076061) were purchased from Sigma Aldrich. Highly modified sgRNA target mouse *TTR* (guide No. G211) was chemically synthesized by AxoLabs based on a previous publication^40^. Modified sgRNA targeting mouse *PCSK9* (cccauaccuuggagcaacgg) and *HPD* (cauucaacgucacaaccacc) were chemically synthesized by Synthego via proprietary specifications.

### mRNA synthesis

ABE8.8 mRNA and CBE4max mRNA were produced by the Drew Weissman laboratory at University of Pennsylvania. Briefly, codon-optimized ABE8.8 or CBE4max sequence was cloned into a proprietary mRNA production plasmid (optimized 3’ and 5’ UTR with a 101 polyA tail), in vitro transcribed in the presence of 1-methyl pseudouridine modified nucleoside, co-transcriptionally capped using the CleanCap technology (TriLink) and cellulose purified to remove double-stranded RNAs. Purified mRNA was ethanol precipitated, washed, resuspended in nuclease-free water and subjected to quality control. All mRNAs were stored at −20 °C until use.

### General procedure for the synthesis of maleates with two different alkyl chains

Aliphatic alcohol (2 mmol, 1 eq.), (Z)-4-((2-ethylhexyl)oxy)-4-oxobut-2-enoic acid (2.2 mmol, 1.1 eq.), DCC (2.2 mmol, 1.1 eq.) and DMAP (0.4 mmol, 0.2 eq.) were dissolved in 20 ml anhydrous DCM in a round bottom flask and stirred at RT under N_2_ protection for 24 h. The reaction mixture was filtered and the filtrate was evaporated under vacuum. The residue was separated using a CombiFlash NextGen 300+ chromatography system (Teledyne Isco) with gradient elution from hexane to 20:80 ethyl acetate/hexane over 15 min to give desired products that were confirmed by ^1^H NMR (**Supplementary Fig. 2**).

### Combinatorial synthesis of ILs

ILs were combinatorially synthesized by solvent-free, catalyst-free Michael addition between amines/thiols and dialkyl maleates. Excess dialkyl maleates (1.25 eq. of –NH_2_/–NH–/–SH) were used to saturate the reaction sites. Taking 5D8 as an example, amine 5 (0.1 mmol, 1 eq.) and D8 (0.25 mmol, 0.25 eq.) were combined in a glass vial and stirred at 80 °C for 24 h. For amine/thiol in the salt form, excess triethylamine was added to neutralize the acid. For insoluble amine/thiol, 100 μL of isopropanol was added to dissolve the reactants. The yield was typically >80%. Crude ILs were dissolved in ethanol and directly used for initial screening. Selected ILs were purified using a CombiFlash NextGen 300+ chromatography system with gradient elution from 100% CH_2_Cl_2_ to 100% CH_2_Cl_2_/MeOH/NH_4_OH (75:22:3) over 10 min to give desired products (**Supplementary Fig. 4**). The lead 5D8 was characterized by mass spectrometry (MS) and nuclear magnetic resonance spectroscopy (^1^H NMR). MS-ESI: calculated for C_48_H_93_N_3_O_8_: 839.70, found [M + H]^+^ = 840.70; ^1^H NMR (600 MHz, CDCl_3_) δ4.05 – 3.75 (m, 10H), 3.52 (q, *J* = 6.2 Hz, 1H), 2.82 – 2.34 (m, 12H), 2.13 (s, 6H), 1.50 (t, *J* = 10.0 Hz, 8H), 1.38 – 1.10 (m, 44H), 0.85 – 0.79 (m, 12H).

### LNP preparation

For initial *in vitro* and *in vivo* screening, LNPs were prepared by pipette mixing of the ethanolic phase containing IL, DOPE, cholesterol and DMG-PEG with the aqueous phase (10 mM citrate buffer, pH 3) containing mLuc at a volume ratio of 1:3. The weight ratio of IL:DOPE:cholesterol:DMG-PEG: mRNA was fixed at 16:10:10:3:1.6. The hydrodynamic size of LNPs formulated by pipette mixing was typically 100-200 nm and the mRNA encapsulation efficiency was typically 60-80%.

For microfluidic formulation of LNPs (including C12-200 LNP), the ethanolic phase containing IL/DOPE/cholesterol/DMG-PEG at a molar ratio of 40:10:48.5:1.5^22^ was mixed with the aqueous phase containing mRNA at a flow rate ratio of 1:3 and at an IL/mRNA weight ratio of 10:1 in a microfluidic device^23^. The benchmark MC3 LNP (or SM-102 LNP) was formulated with MC3 (or SM-102), DSPC, cholesterol and DMG-PEG at a molar ratio of 50:10:38.5:1.5 using microfluidic mixing at an ionizable lipid/mRNA weight ratio of 10:1. The LP-01 LNP was formulated similarly according to a previous study^5^. LNPs were dialyzed against 1×PBS in a 20 kDa MWCO cassette for 2 h, filtered through a 0.22 μM filter and stored at 4 °C. The hydrodynamic size of LNPs formulated by microfluidic mixing was typically 80-120 nm and the mRNA encapsulation efficiency was typically >90%.

### Characterization

^1^H-NMR was recorded using a Bruker 400 MHz NMR spectrometer. MS was performed on a Waters Acquity LC-MS system equipped with UV-Vis and MS detectors. The hydrodynamic size, polydispersity index (PDI) and zeta potential of LNPs were measured using a Malvern Zetasizer Nano ZS90. The morphology of LNPs was characterized by a cryo-electron microscope (Titan Krios, Thermo Fisher) equipped with a K3 Bioquantum. The mRNA encapsulation efficiency and the p*K*_a_ of LNP were determined using a modified Quant-iT RiboGreen RNA assay (Invitrogen) and a 6-(p-toluidinyl)naphthalene-2-sulfonic acid (TNS) assay^24^, respectively.

### Cell culture and animal studies

Human hepatocellular carcinoma HepG2 cells were purchased from American Type Culture Collection (ATCC) and maintained in Dulbecco’s Modified Eagle Medium (DMEM) supplemented with 10% fetal bovine serum (FBS), 100 U/mL penicillin and 100 μg/mL streptomycin. Cells were cultured at 37 °C in a humidified incubator of 5% CO_2_, and routinely tested for mycoplasma contamination.

All animal protocols were approved by the Institutional Animal Care and Use Committee (IACUC) of University of Pennsylvania (#806540), and animal procedures were performed in accordance with the Guidelines for Care and Use of Laboratory Animals at the University of Pennsylvania. C57BL/6 female mice (6-8 weeks, 18-20 g) were purchased from The Jackson Laboratory. *FAH*^−/−^ male mice (6-8 months, 25-30 g) were bred in our animal facility and genotyped using a previously described protocol^21^. They were maintained on nitisinone (Yecuris Corporation) in their drinking water at a concentration of 16.5 mg/L. Mice were housed in a specific-pathogen-free animal facility at ambient temperature (22 ± 2 °C), air humidity 40%–70% and 12-h dark/12-h light cycle.

### *In vitro* LNP screening

For high-throughput *in vitro* screening of LNPs, HepG2 cells were seeded in 384-well plates at a density of 1,000 per well overnight and mLuc-loaded LNPs were used to treat cells at an mRNA dose of 3 ng/well for 24 h. For *in vitro* batch screening of LNPs, HepG2 cells were seeded in 96-well plates at a density of 5,000 per well overnight. mLuc-LNPs with the same amine structure were pooled and used to treat cells at an mRNA dose of 15 ng/well for 24 h. Luciferase expression was evaluated by Luciferase Reporter 1000 Assay System (Promega, #E4550) according to the manufacturer’s protocol.

### *In vivo* LNP screening

For *in vivo* screening of LNPs, mice were i.v. injected with either pooled LNPs or individual LNPs at an mLuc dose of 0.1 mg/kg. Four hours later, mice were intraperitoneally injected with D-luciferin potassium salt (150 mg/kg), and bioluminescence imaging was performed using an *in vivo* imaging system (PerkinElmer). Whole-body total flux was quantified and normalized to C12-200 LNP-treated mice.

### Isolation and identification of plasma proteins absorbed to LNPs

Plasma proteins absorbed to LNPs were isolated as previously described. Briefly, mouse blood was collected into EDTA-treated tubes and centrifuged at 3,000 × *g* at 4 °C for 10 min to obtain plasma. Plasma was further centrifuged at 13,000 × *g* at 4 °C for 30 min to remove protein aggregates. LNPs were mixed with an equal volume of plasma at a final mRNA concentration of 0.1 mg/mL and incubated for 1 h at 37 °C under shaking. The plasma protein-coated LNPs were isolated by centrifugation at 13,000× *g* at 4 °C for 30 min, followed by washing with cold PBS three times to remove unbound proteins. The protein concentration was determined using a BCA method before protein analysis by the Proteomics and Metabolomics Core at the Wistar Institute.

### Systemic delivery of CRISPR gene editor targeting TTR

C57BL/6 mice were i.v. injected with Cas9 mRNA/TTR sgRNA (4:1, wt:wt)-loaded LNPs at a total RNA dose of 1 mg/kg. Serum was collected on day 7 and analyzed by ELISA (Aviva Systems Biology, #OKIA00111). Mice were euthanized, and livers were collected to determine the on-target indel frequency by next-generation sequencing (NGS). For TTR on-target DNA sequencing, DNA was extracted from the liver using the Qiagen Puregene Tissue Kit (#158063) and quantified using a Nanodrop 2000. PCR amplification of the TTR target site was carried out using Q5 High-Fidelity DNA Polymerase (#M0491, New England Biolabs) and the following primers: 5’-CGGTTTACTCTGACCCATTTC-3’ and 5’-GGGCTTTCTACAAGCTTACC-3’. Deep sequencing of the TTR amplicons and determination of the on-target indel frequency was performed essentially as described except that 150 bp pair end reads were produced^25^.

### Systemic delivery of ABE base editor targeting PCSK9

C57BL/6 mice were i.v. injected with ABE8.8 mRNA/PCSK9 sgRNA (4:1, wt:wt)-loaded LNPs at a total RNA dose of 0.75 mg/kg. Serum was collected on day 7 and analyzed by ELISA (#ab215538, Abcam). Mice were euthanized, and livers were collected. Genomic DNA was extracted for the analysis of on-target *PCSK9* editing by NGS. PCR amplification of the PCSK9 target site was carried out with the following primers: 5’-GGCTGCACTTAGAGACCACC-3’ and 5’-ATGAAGAGCTGATGCTCGCC-3’. Deep sequencing of the PCSK9 amplicons was performed.

### Systemic delivery of CBE base editor targeting HPD

*FAH*^−/−^ mice maintained on nitisinone were i.v. injected with CBE4max mRNA/HPD sgRNA (4:1, wt:wt)-loaded LNPs at a total RNA dose of 0.6 mg/kg. On day 7, mice were euthanized, and livers were collected. Genomic DNA was extracted for the analysis of on-target *HPD* editing by NGS. PCR amplification of the HPD target site was carried out with the following primers: 5’-CCTTCCTTTAACAGAGCCCACT-3’ and 5’-TGGGTAAGATTTCGCAGGCA-3’. Deep sequencing of the HPD amplicons was performed. For the survival study, nitisinone was withdrawn two weeks post-treatment and the survival of *FAH*^−/−^ mice was monitored.

### Systemic delivery of siRNA targeting TTR

Three TTR siRNAs were pooled at a 1:1:1 molar ratio and encapsulated into LNPs using microfluidic mixing. Mice were i.v. injected with TTR siRNA-loaded LNPs at a total siRNA dose of 0.05 mg/kg. Serum was collected on day 3 and analyzed by ELISA.

### Liver toxicity evaluation

Serum was collected at 24 h post injection of Cas9 mRNA/TTR sgRNA-loaded LNPs at a total RNA dose of 1 mg/kg. ALT and AST activities were determined by alanine transaminase colorimetric activity assay kit (#700260, Cayman) and aspartate aminotransferase colorimetric activity assay kit (#701640, Cayman), respectively.

### Statistical analysis

Data are presented as mean ± SD. Student’s *t*-test or one-way analysis of variance (ANOVA) followed by Tukey’s test was applied for comparison between two groups or among multiple groups using Graphpad Prism 8.0, respectively. *p* < 0.05 was considered to be statistically significant.

## Data availability

All relevant data supporting the findings of this study are available within the paper and the Supplementary Information.

## Acknowledgments

MJM acknowledges support from a US National Institutes of Health (NIH) Director’s New Innovator Award (DP2 TR002776), a Burroughs Wellcome Fund Career Award at the Scientific Interface (CASI), a US National Science Foundation CAREER Award (CBET-2145491), and an American Cancer Society Research Scholar Grant (RSG-22-122-01-ET). JMW acknowledges support from iECURE. The authors acknowledge Dr. Stefan Steimle from the Beckman Center for Cryo Electron Microscopy at UPenn Perelman School of Medicine (RRID: SCR_022375) for the help in characterizing the morphology of LNPs. The authors acknowledge UPenn Gene Therapy Program NAT Core for sequencing service and thank Kelly Martins for processing the NGS data. The authors acknowledge Wistar Institute’s Proteomics and Metabolomics Core for plasma protein analysis.

## Author Contributions

Conceptualization: XH, YX, MJM; Methodology: XH, YX, AR, JX; Investigation: XH, YX, AR, JX, RP, VC; Visualization: XH, YX; Funding acquisition: JMW, DW, MJM; Supervision: JMW, DW, MJM; Writing – original draft: XH, YX, AR, RP, MJM; Writing – review & editing: YX, AR, JX, RP, LX, NG, VC, MGA, JMW, DW, MJM.

