## Supplementary Information for "Plug-and-play assembly of biodegradable ionizable lipids for potent mRNA delivery and gene editing *in vivo*"

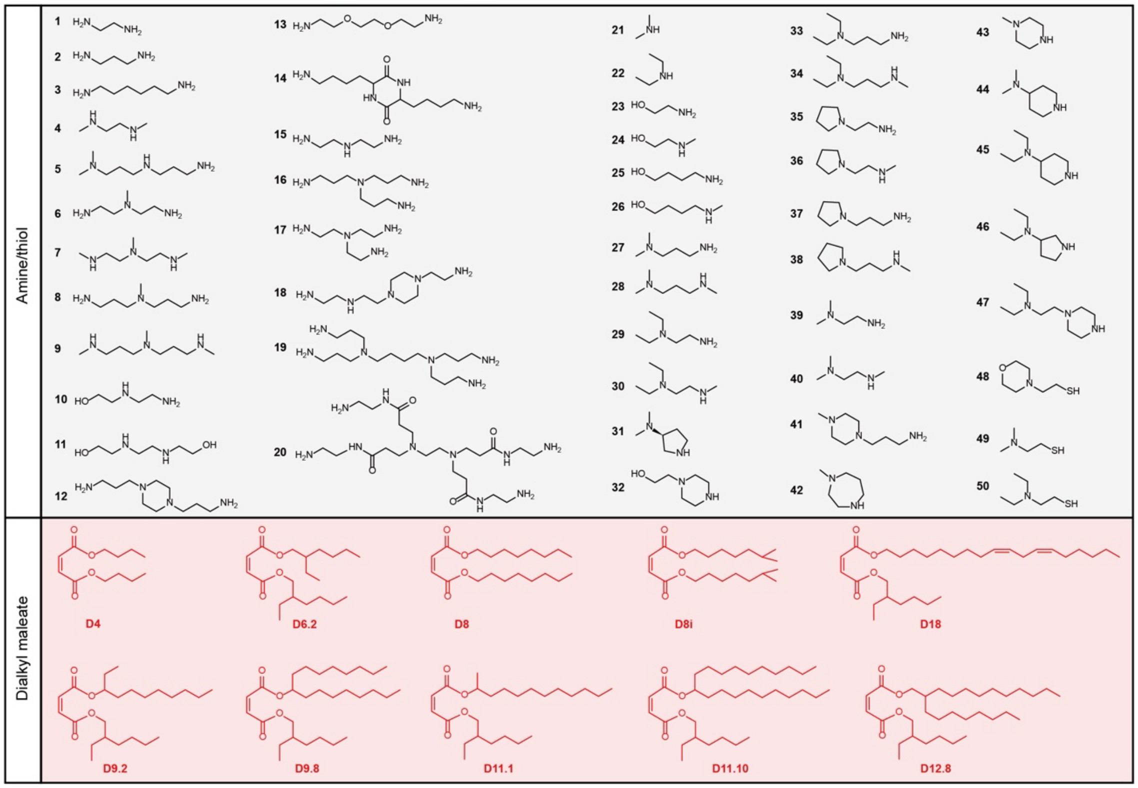


**Supplementary Fig. 1** Amines/thiols and dialkyl maleates used in the library.

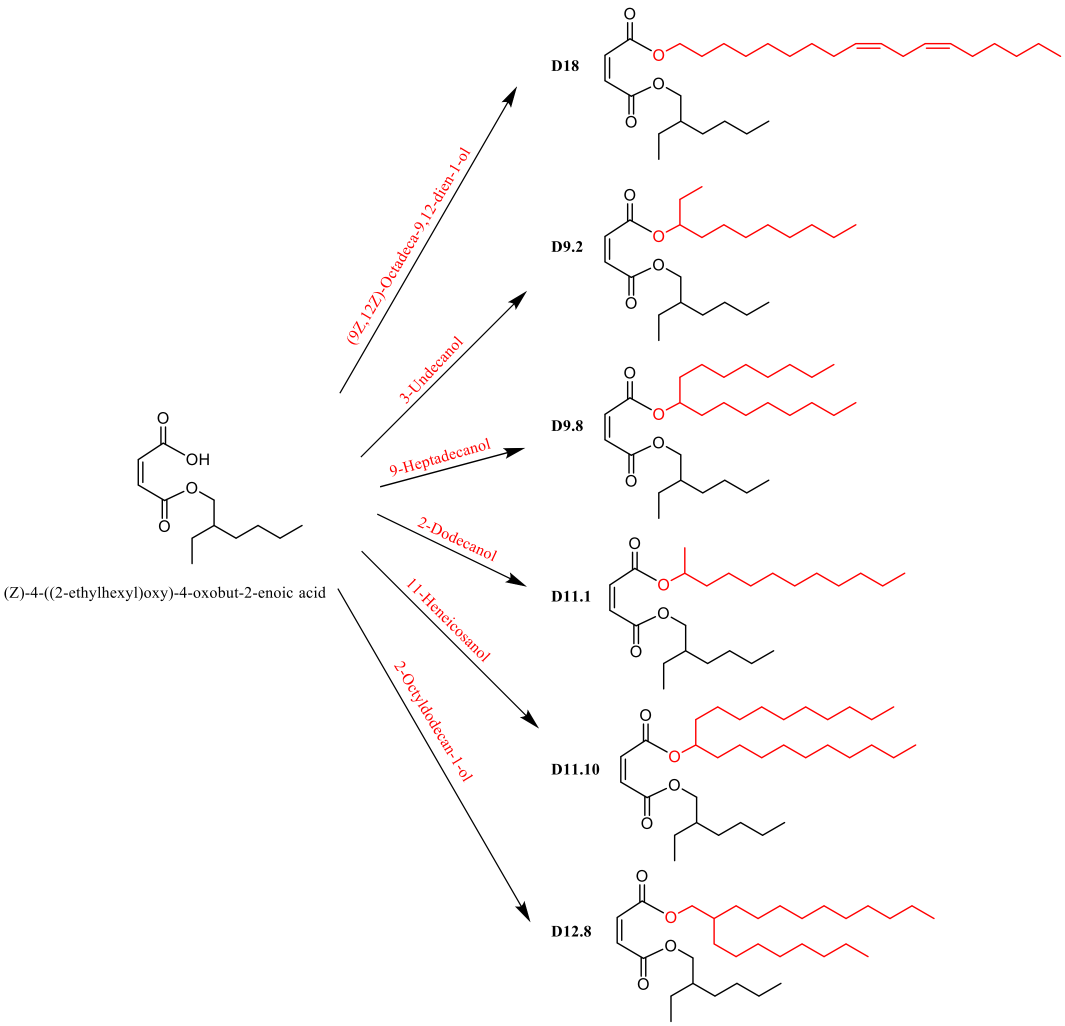


**Supplementary Fig. 2** Synthetic routes for maleates with two different alkyl chains. D18, D9.2, D9.8, D11.1, D11.10 and D12.8 were synthesized via a DCC/DMAP-mediated esterification reaction.

**D18**: Light yellow oil, yield 92%. $^1H$ NMR (400 MHz, $CDCl_3$) δ 6.23 (s, 2H), 5.44 – 5.27 (m, 4H), 4.20 – 4.08 (m, 4H), 2.77 (t, J = 6.5 Hz, 2H), 2.10 – 2.00 (m, 4H), 1.70 – 1.58 (m, 1H), 1.43 – 1.24 (m, 26H), 0.89 (td, J = 7.2, 3.8 Hz, 9H).

**D9.2**: Colorless oil, yield 90%. $^1H$ NMR (400 MHz, $CDCl_3$) δ 6.30 – 6.18 (m, 2H), 4.13 (ddd, J = 10.7, 5.9, 3.7 Hz, 3H), 1.67 – 1.61 (m, 1H), 1.45 – 1.24 (m, 24H), 0.96 – 0.87 (m, 12H).

**D9.8**: Colorless oil, yield 88%. $^1H$ NMR (400 MHz, $CDCl_3$) δ 6.30 – 6.17 (m, 2H), 4.13 (dddd, J = 10.2, 6.1, 4.3, 1.3 Hz, 3H), 1.58 (s, 1H), 1.46 – 1.23 (m, 36H), 0.95 – 0.87 (m, 12H).

**D11.1**: Colorless oil, yield 90%. $^1H$ NMR (400 MHz, $CDCl_3$) δ 6.23 (d, J = 2.6 Hz, 2H),

4.18 – 4.06 (m, 3H), 1.86 – 1.59 (m, 1H), 1.46 – 1.21 (m, 29H), 0.91 (td, J = 7.3, 5.3 Hz, 9H).

**D11.10**: Colorless oil, yield 85%. $^{1}H$ NMR (400 MHz, $CDCl_3$) δ 6.30 – 6.16 (m, 2H), 4.18 – 4.08 (m, 3H), 1.65 (s, 1H), 1.49 – 1.22 (m, 44H), 0.97 – 0.86 (m, 12H).

**D12.8**: Colorless oil, yield 85%. $^{1}H$ NMR (400 MHz, $CDCl_3$) δ 6.25 (s, 2H), 4.20 – 4.06 (m, 4H), 1.72 – 1.62 (m, 2H), 1.47 – 1.24 (m, 40H), 0.91 (td, J = 7.2, 4.9 Hz, 12H).

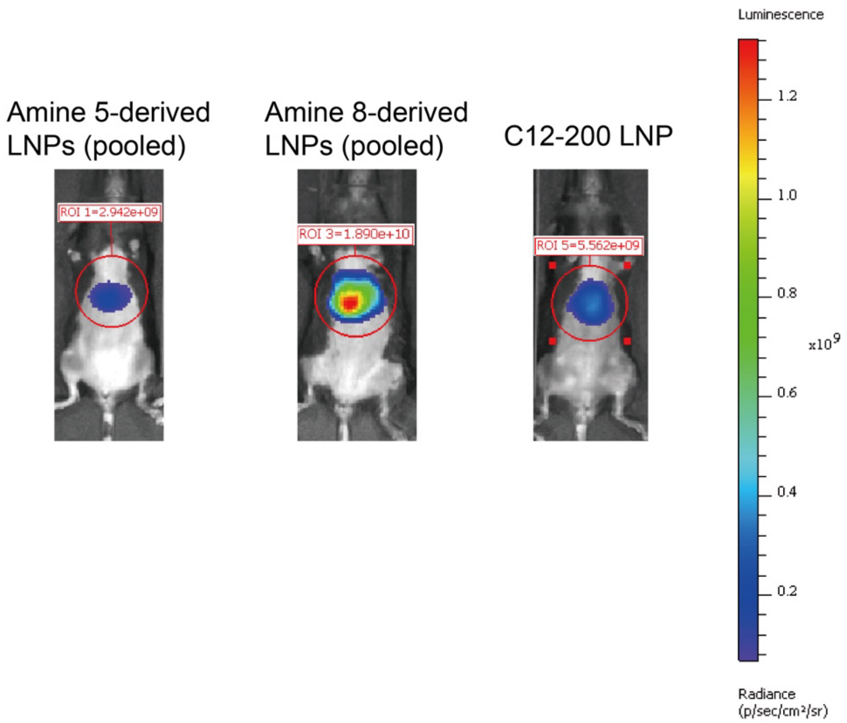


**Supplementary Fig. 3** Representative *in vivo* bioluminescence imaging results of batch LNP screening. mLuc-LNPs with the same amine structure were pooled and i.v. injected into mice at an mRNA dose of 0.1 mg/kg. Images were taken at 4 h post-treatment.

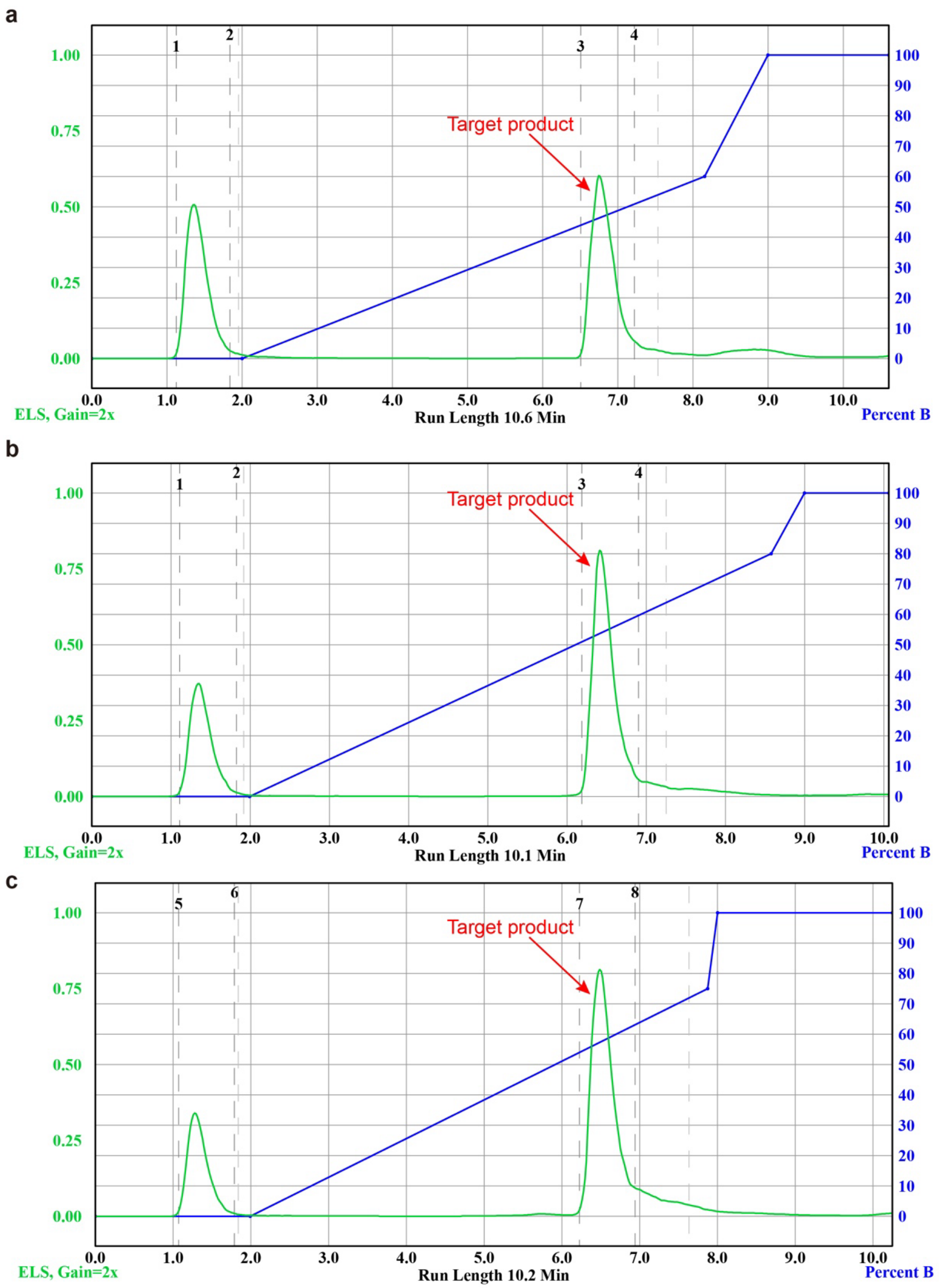


**Supplementary Fig. 4** Representative purification traces of ILs. **a**, 5D8. **b**, 8D6.2. **c**, 12D6.2. The purification trace was recorded by a CombiFlash NextGen 300+ chromatography system equipped with an evaporative light scattering (ELS) detector. The crude product was purified with gradient elution from 100% $CH_2Cl_2$ to 100% $CH_2Cl_2$/MeOH/$NH_4OH$ (75:22:3) over 10 min to give target product.

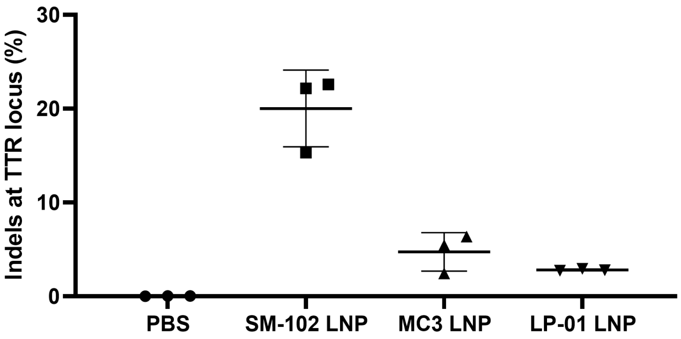


**Supplementary Fig. 5** Indels at TTR locus. Mice were i.v. injected with LNPs encapsulating Cas9 mRNA/TTR sgRNA (4:1, wt:wt) at a total RNA dose of 1 mg/kg. On day 7, DNA was extracted from the liver to determine on-target indel frequency by next-generation sequencing. Data are presented as mean ± SD (n=3 independent biological replicates).

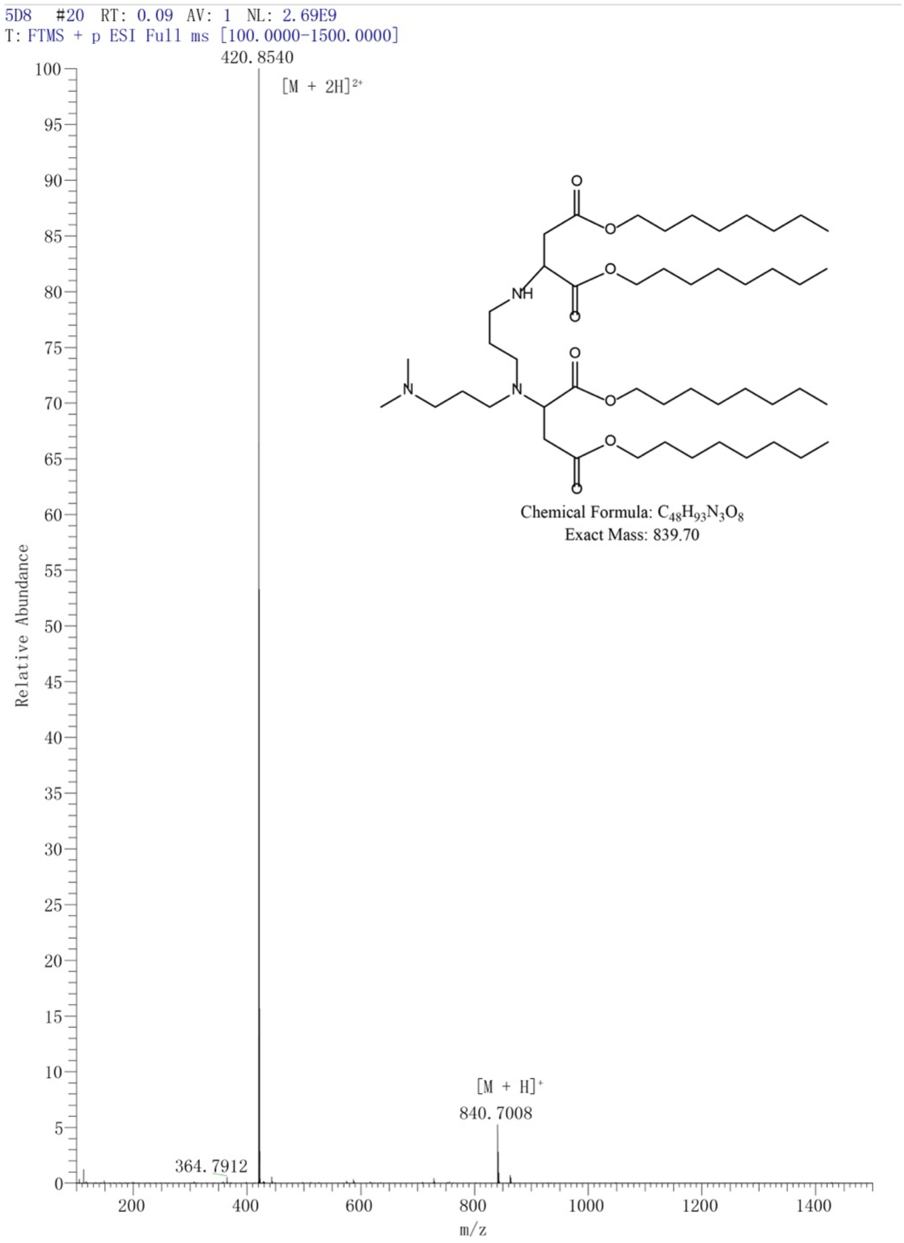


**Supplementary Fig. 6** Mass spectrum of 5D8.

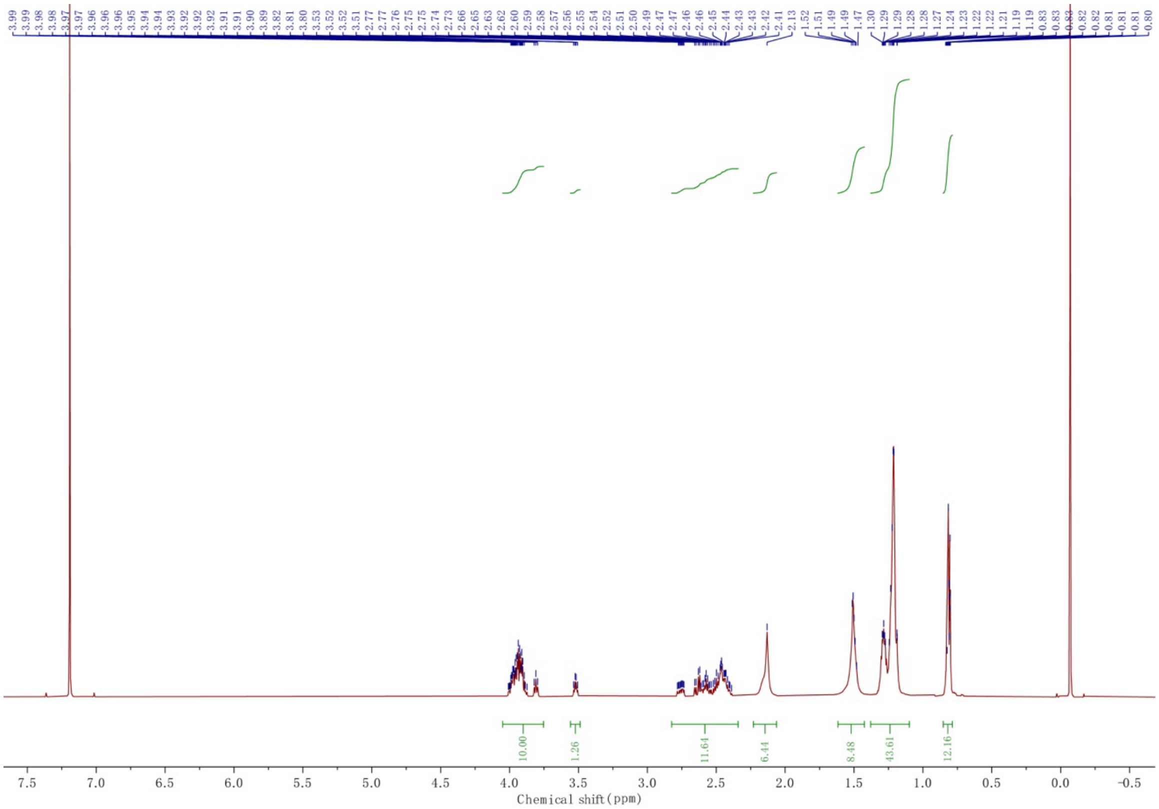

**Supplementary Fig. 7** $^{1}H$ NMR spectrum of 5D8 in $CDCl_3$.

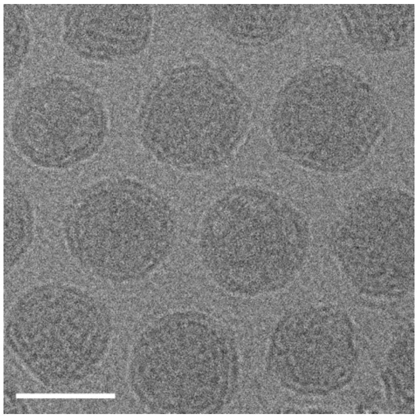

**Supplementary Fig. 8** A representative cryo-EM image of 5D8 LNP from three independent experiments. Scale bar = 50 nm.

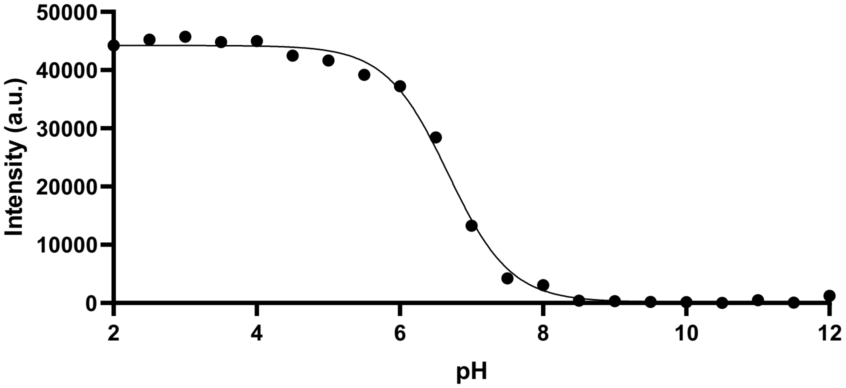


**Supplementary Fig. 9** TNS assay was used to determine the apparent p$K_a$ of 5D8 LNP. TNS fluorescence signal corresponds to ionization. p$K_a$ is calculated as the pH corresponding to half of the maximum TNS fluorescence value.

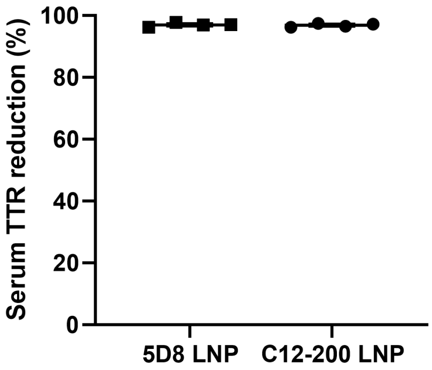


**Supplementary Fig. 10** LNP-mediated TTR siRNA delivery *in vivo*. Mice were i.v. injected with TTR siRNA-loaded LNPs at a dose of 0.05 mg/kg. Serum was collected on day 3 for ELISA analysis. Both 5D8 and C12-200 LNPs achieved ~100% reduction of serum TTR. Data are presented as mean ± SD (n=4 independent biological replicates).

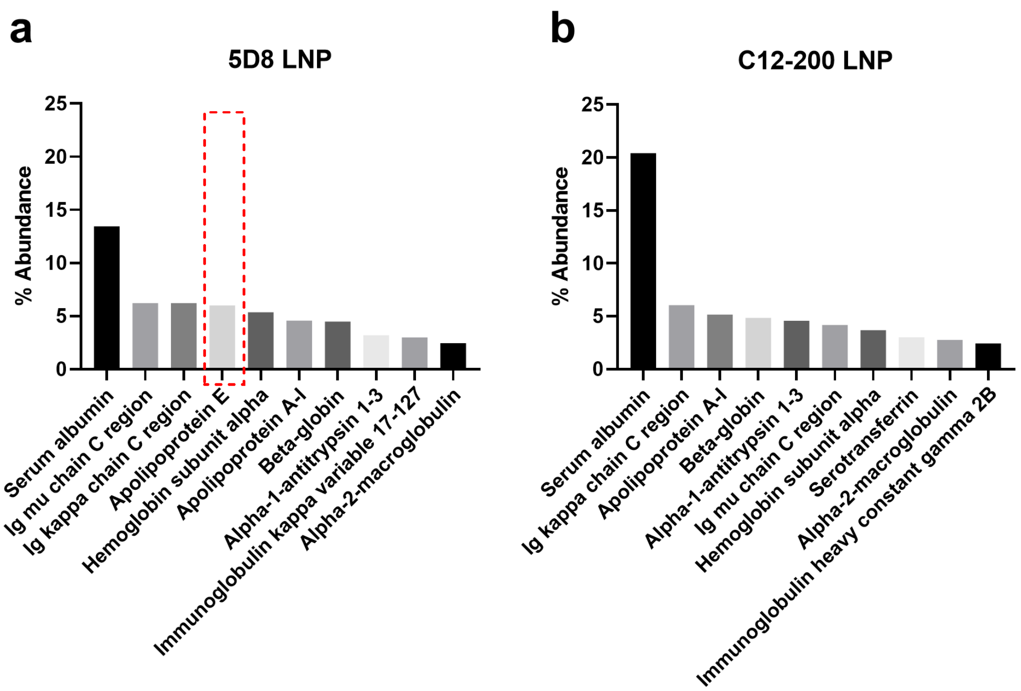


**Supplementary Fig. 11** The top 10 most abundant plasma proteins that bind LNPs. **a**, 5D8 LNP. **b**, C12-200 LNP. Apolipoprotein E (highlighted in the dashed red rectangle) is highly enriched on the surface of 5D8 LNP compared to C12-200 LNP.

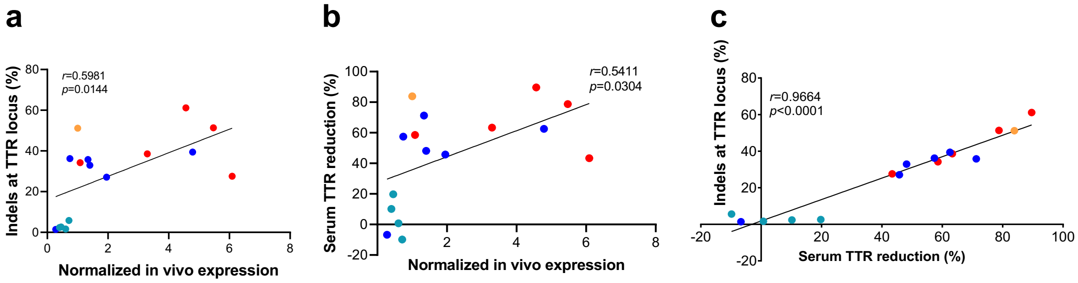


**Supplementary Fig. 12** Correlation analysis. **a**, Correlation between in vivo mLuc delivery efficiency and TTR gene editing efficiency. Statistical significance was evaluated by a two-tailed correlation analysis using GraphPad Prism 8.0. **b**, Correlation between in vivo mLuc delivery efficiency and serum TTR reduction. Statistical significance was evaluated by a two-tailed correlation analysis using GraphPad Prism 8.0. **c**, Correlation between serum TTR reduction and TTR gene editing efficiency. Statistical significance was evaluated by a two-tailed correlation analysis using GraphPad Prism 8.0. The red dots indicate amine 5-derived ILs. The blue dots indicate amine 8-derived ILs. The cyan dots indicate amine 12-derived ILs. The brown dot indicates C12-200.

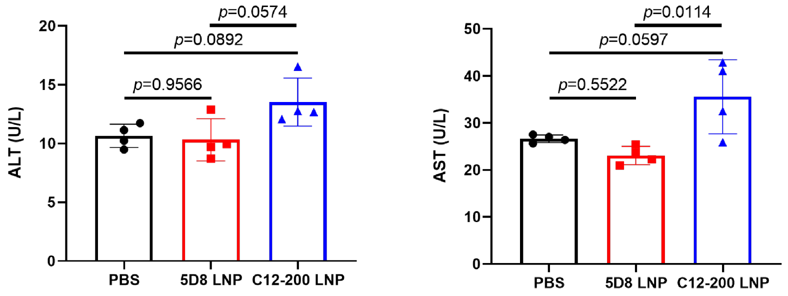


**Supplementary Fig. 13** ALT and AST analysis. LNPs encapsulating Cas9 mRNA/TTR sgRNA (4:1, wt:wt) were i.v. injected into mice at a total RNA dose of 1 mg/kg. Serum was collected for ALT and AST analysis at 24 h post-treatment. Data are presented as mean ± SD (n=4 independent biological replicates). Statistical significance was evaluated by a one-way ANOVA with Tukey's correction.

**Supplementary Table 1** Physiochemical parameters of 5D8 LNP and C12-200 LNP formulated by microfluidic mixing.

| **LNP** | **Size (nm)** | **PDI** | **Zeta potential (mV)** | **EE (%)** | **p*K*a** |
|---|---|---|---|---|---|
| 5D8 | 87.2 ± 2.4 | 0.144 | 0.71 ± 0.91 | 96.9 ± 1.5 | 6.58 |
| C12-200 | 101.8 ± 0.8 | 0.142 | -1.82 ± 0.60 | 93.1 ± 1.7 | 6.76 |

The hydrodynamic size, PDI and zeta potential of LNPs were obtained by dynamic light scattering (DLS) measurement in PBS (pH 7.4). The mRNA encapsulation efficiency (EE) was determined using a modified Quant-iT RiboGreen RNA assay. The p$K_a$ of LNP were determined using a 6-(p-toluidinyl)naphthalene-2-sulfonic acid (TNS) assay. Data are presented as mean ± SD (n=3 independent biological replicates).
